## Supporting information for "A deep dive into VDAC1 conformational diversity using all-atom simulations provides new insights into the structural origin of the closed states"

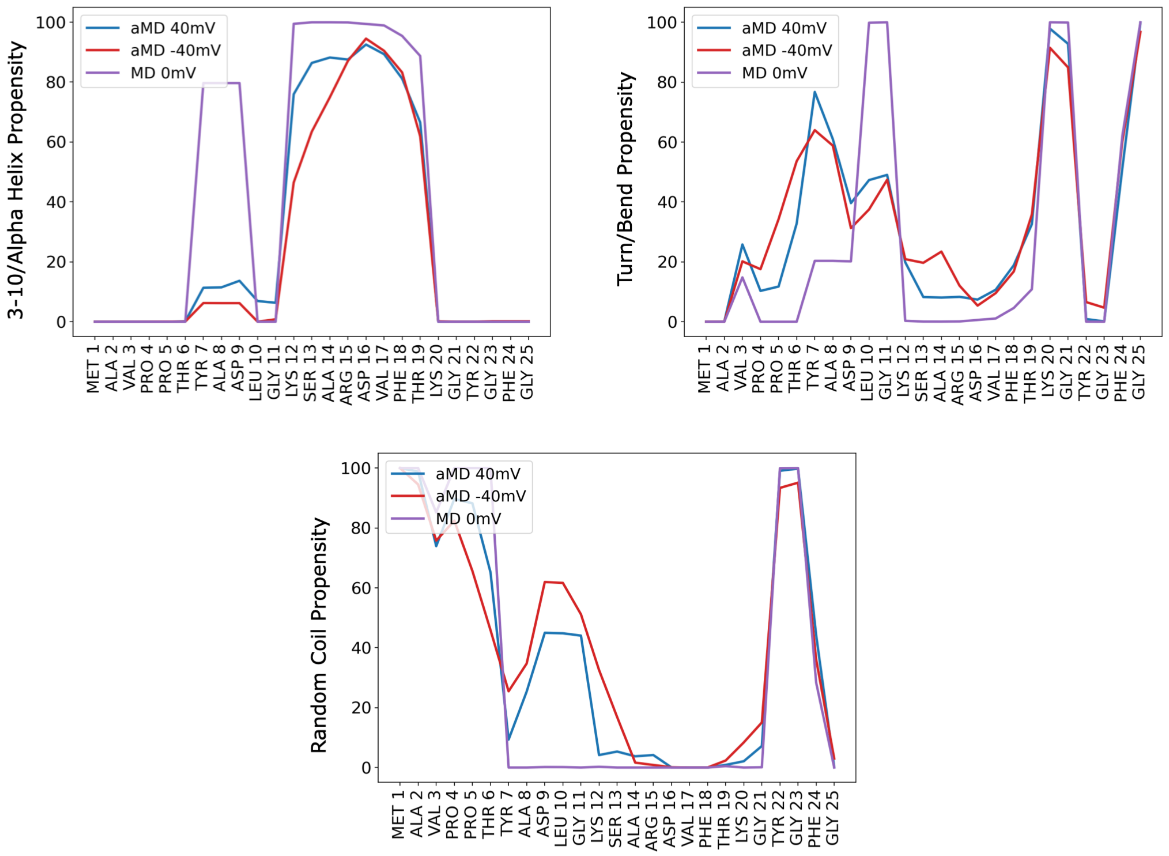


**Figure S1.** Secondary structure propensities obtained from exploratory aMD simulations at applied voltage. As a comparison, propensities obtained from unbiased MD at 0 mV, starting from the 3EMN structure, were also displayed. Notably, the unbiased MD trajectory did not reveal any major conformational change with respect to the crystal structure. Propensities of helices ($3_{10}$- and alpha), turn/bends as well as random coiled are shown.

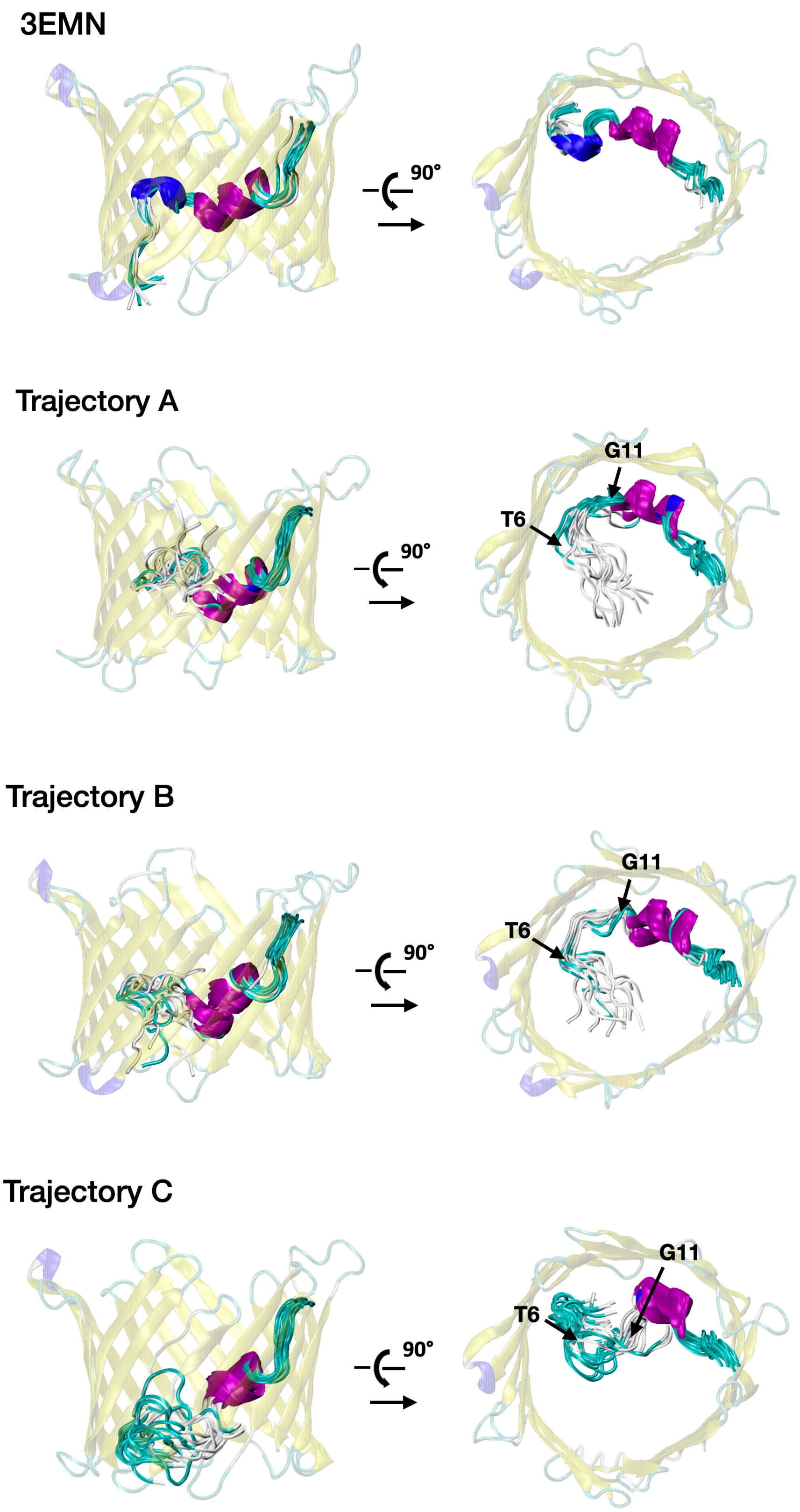


**Figure S2.** Frames extracted from unbiased MD trajectories at applied voltage. Trajectories are 500 ns-long and include the one initiated from 3EMN as well as trajectories A, B and C highlighted in the main text. On each figure, 13 frames were taken at regular time interval and superimposed. Only fluctuations in the N-terminal region are shown.

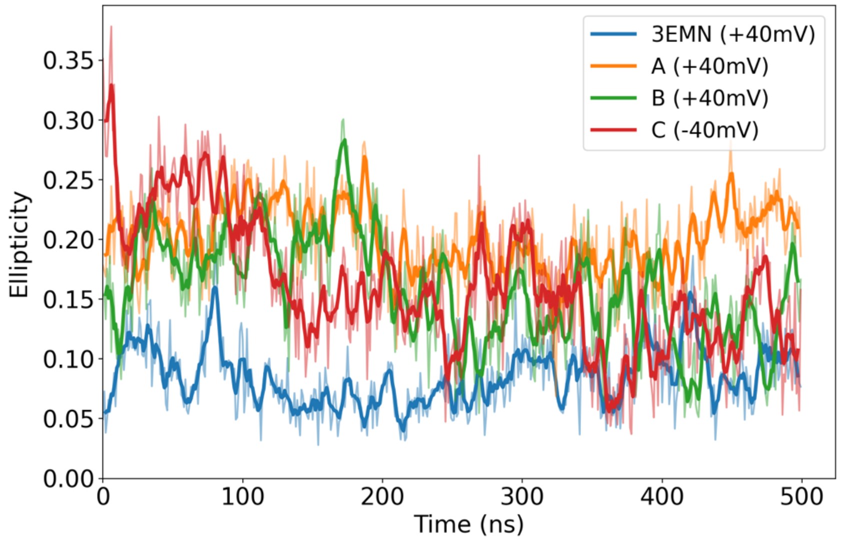


**Figure S3.** Ellipticity measured during unbiased MD at applied voltage. Simulations were initiated from 3EMN and structures A, B and C.

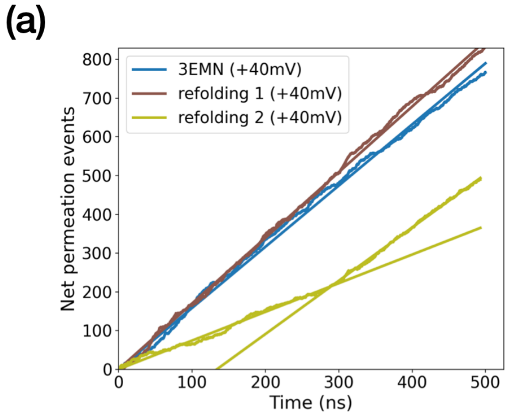


(b)

| Trajectory | Conductance (nS) | % open state |
|---|---|---|
| 3EMN | 3.71 | 100.0 % |
| refolding 1 | 3.98 | 107.3 % |
| refolding 2 (t < 270 ns) | 1.74 | 46.8 % |
| refolding 2 (t > 270ns) | 3.23 | 87.1 % |

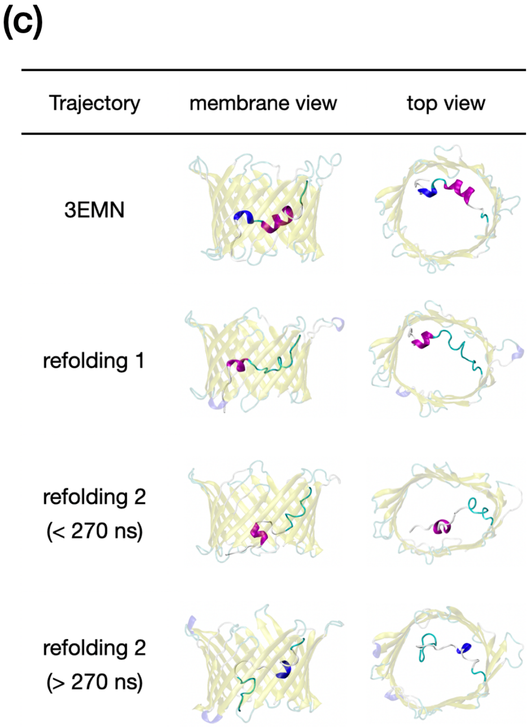


**Figure S4. (a)** Net ion permeation events recorded from unbiased MD at applied voltage. Trajectories were initiated from 2 structures exhibiting a refolded N-terminal domain. For each curve, simple linear regression (straight lines plotted along with each curve) was performed. As refolding 2 exhibited two conducting regimes, two linear regressions, one before and one after the bifurcation point were done. Bifurcation points for refolding 2 was estimated at 270 ns. **(b)** Conductance deduced from the slope of the ion permeation curves. The slope corresponds to the electric current, which after renormalization and division by the applied voltage provides an estimate of the conductance. **(c)** Average structures of MD trajectories in side view (membrane) and top view.

**Table S1.** GCMC/BD results obtained at 1M KCl on 200 frames extracted from aMD simulations at +40mV. For each frame, the conductance $\sigma = I/V$ and the current ratio $I_{Cl-}/I_{K+}$ are provided. Frames highlighted in orange (55 frames) show a conductance less than 2.5 nS. These were used as starting point in MD simulations to better evaluate their conductance and stability.

| # | $\sigma$ | $I_{Cl-}/I_{K+}$ | # | $\sigma$ | $I_{Cl-}/I_{K+}$ | # | $\sigma$ | $I_{Cl-}/I_{K+}$ | # | $\sigma$ | $I_{Cl-}/I_{K+}$ |
|---|---|---|---|---|---|---|---|---|---|---|---|
| 1 | 0.52 | 1.60 | 51 | 2.42 | 2.56 | 101 | 2.86 | 2.58 | 151 | 3.38 | 5.03 |
| 2 | 0.70 | 7.75 | 52 | 2.42 | 3.65 | 102 | 2.88 | 1.25 | 152 | 3.41 | 1.74 |
| 3 | 1.10 | 5.88 | 53 | 2.42 | 10.00 | 103 | 2.90 | 2.15 | 153 | 3.42 | 3.17 |
| 4 | 1.10 | 5.11 | 54 | 2.48 | 1.76 | 104 | 2.90 | 2.09 | 154 | 3.42 | 3.07 |
| 5 | 1.14 | 18.00 | 55 | 2.48 | 2.54 | 105 | 2.94 | 2.00 | 155 | 3.42 | 1.67 |
| 6 | 1.18 | ¥ | 56 | 2.50 | 2.38 | 106 | 2.94 | 2.20 | 156 | 3.42 | 2.23 |
| 7 | 1.32 | 3.40 | 57 | 2.50 | 1.27 | 107 | 2.96 | 4.48 | 157 | 3.43 | 2.49 |
| 8 | 1.32 | 65.01 | 58 | 2.50 | 5.94 | 108 | 2.98 | 3.38 | 158 | 3.43 | 1.76 |
| 9 | 1.40 | 1.69 | 59 | 2.52 | 1.52 | 109 | 2.98 | 2.04 | 159 | 3.43 | 1.55 |
| 10 | 1.42 | 1.63 | 60 | 2.52 | 10.45 | 110 | 3.00 | 4.00 | 160 | 3.43 | 7.15 |
| 11 | 1.56 | 1.89 | 61 | 2.58 | 1.75 | 111 | 3.00 | 2.33 | 161 | 3.46 | 3.44 |
| 12 | 1.56 | 2.90 | 62 | 2.58 | 2.58 | 112 | 3.02 | 5.86 | 162 | 3.46 | 1.54 |
| 13 | 1.58 | 6.90 | 63 | 2.58 | 2.79 | 113 | 3.02 | 2.51 | 163 | 3.46 | 2.15 |
| 14 | 1.66 | 4.93 | 64 | 2.58 | 2.69 | 114 | 3.04 | 2.23 | 164 | 3.46 | 2.15 |
| 15 | 1.66 | 4.19 | 65 | 2.58 | 3.96 | 115 | 3.04 | 2.80 | 165 | 3.50 | 3.86 |
| 16 | 1.68 | 3.00 | 66 | 2.60 | 1.89 | 116 | 3.04 | 1.34 | 166 | 3.51 | 2.43 |
| 17 | 1.78 | 11.72 | 67 | 2.60 | 6.22 | 117 | 3.04 | 1.87 | 167 | 3.53 | 1.10 |
| 18 | 1.82 | 3.14 | 68 | 2.62 | 1.18 | 118 | 3.04 | 1.92 | 168 | 3.55 | 1.60 |
| 19 | 1.82 | 2.25 | 69 | 2.62 | 2.85 | 119 | 3.04 | 2.53 | 169 | 3.55 | 1.77 |
| 20 | 1.82 | 2.79 | 70 | 2.62 | 1.85 | 120 | 3.06 | 6.65 | 170 | 3.55 | 4.37 |
| 21 | 1.84 | 3.00 | 71 | 2.62 | 3.23 | 121 | 3.06 | 2.40 | 171 | 3.59 | 2.09 |
| 22 | 1.86 | 2.32 | 72 | 2.66 | 3.75 | 122 | 3.12 | 9.39 | 172 | 3.61 | 1.73 |
| 23 | 1.88 | 1.85 | 73 | 2.66 | 1.83 | 123 | 3.14 | 2.02 | 173 | 3.63 | 4.33 |
| 24 | 1.88 | 7.55 | 74 | 2.66 | 4.32 | 124 | 3.14 | 3.24 | 174 | 3.67 | 1.35 |
| 25 | 1.90 | 1.71 | 75 | 2.68 | 3.32 | 125 | 3.14 | 1.53 | 175 | 3.69 | 3.60 |
| 26 | 1.98 | 4.21 | 76 | 2.70 | 2.07 | 126 | 3.18 | 3.82 | 176 | 3.71 | 1.98 |
| 27 | 2.00 | 4.26 | 77 | 2.70 | 2.14 | 127 | 3.18 | 1.37 | 177 | 3.72 | 2.32 |
| 28 | 2.02 | 4.32 | 78 | 2.72 | 1.72 | 128 | 3.18 | 1.89 | 178 | 3.75 | 4.06 |
| 29 | 2.06 | 1.29 | 79 | 2.72 | 2.78 | 129 | 3.20 | 2.08 | 179 | 3.78 | 2.57 |
| 30 | 2.08 | 5.50 | 80 | 2.74 | 5.52 | 130 | 3.20 | 2.48 | 180 | 3.78 | 2.26 |

| | | | | | | | | | | | |
|---|---|---|---|---|---|---|---|---|---|---|---|
| 31 | 2.08 | 5.12 | 81 | 2.74 | 5.23 | 131 | 3.20 | 1.42 | 181 | 3.82 | 2.29 |
| 32 | 2.12 | 3.24 | 82 | 2.74 | 3.72 | 132 | 3.20 | 2.56 | 182 | 3.85 | 3.37 |
| 33 | 2.14 | 2.34 | 83 | 2.74 | 8.79 | 133 | 3.20 | 1.91 | 183 | 3.86 | 2.71 |
| 34 | 2.14 | 4.10 | 84 | 2.74 | 1.02 | 134 | 3.20 | 3.21 | 184 | 3.87 | 1.80 |
| 35 | 2.16 | 2.48 | 85 | 2.74 | 2.70 | 135 | 3.22 | 3.88 | 185 | 3.87 | 1.80 |
| 36 | 2.24 | 9.18 | 86 | 2.74 | 2.91 | 136 | 3.24 | 4.06 | 186 | 3.87 | 2.94 |
| 37 | 2.28 | 2.35 | 87 | 2.76 | 1.71 | 137 | 3.26 | 5.04 | 187 | 3.91 | 3.54 |
| 38 | 2.28 | 3.96 | 88 | 2.76 | 6.67 | 138 | 3.26 | 2.40 | 188 | 3.95 | 3.58 |
| 39 | 2.32 | 0.97 | 89 | 2.76 | 4.11 | 139 | 3.28 | 4.47 | 189 | 3.97 | 1.61 |
| 40 | 2.32 | 2.05 | 90 | 2.78 | 1.84 | 140 | 3.28 | 3.82 | 190 | 4.00 | 2.12 |
| 41 | 2.32 | 1.15 | 91 | 2.78 | 4.35 | 141 | 3.28 | 1.83 | 191 | 4.01 | 3.35 |
| 42 | 2.34 | 1.54 | 92 | 2.78 | 2.97 | 142 | 3.28 | 1.73 | 192 | 4.05 | 2.74 |
| 43 | 2.34 | 1.85 | 93 | 2.78 | 3.09 | 143 | 3.30 | 1.85 | 193 | 4.08 | 2.04 |
| 44 | 2.34 | 2.34 | 94 | 2.80 | 1.80 | 144 | 3.30 | 1.75 | 194 | 4.08 | 2.14 |
| 45 | 2.36 | 2.03 | 95 | 2.80 | 3.38 | 145 | 3.30 | 1.66 | 195 | 4.09 | 2.92 |
| 46 | 2.36 | 7.43 | 96 | 2.80 | 1.55 | 146 | 3.32 | 4.53 | 196 | 4.19 | 2.67 |
| 47 | 2.38 | 4.18 | 97 | 2.82 | 14.66 | 147 | 3.32 | 2.26 | 197 | 4.21 | 1.53 |
| 48 | 2.38 | 2.13 | 98 | 2.82 | 2.20 | 148 | 3.34 | 3.77 | 198 | 4.21 | 2.33 |
| 49 | 2.40 | 1.35 | 99 | 2.82 | 3.70 | 149 | 3.35 | 2.15 | 199 | 4.33 | 3.70 |
| 50 | 2.40 | 1.79 | 100 | 2.86 | 4.30 | 150 | 3.36 | 5.00 | 200 | 4.39 | 3.87 |

**Table S2.** GCMC/BD results obtained at 1M KCl on 200 frames extracted from aMD simulations at -40mV. For each frame, the conductance $\sigma = I/V$ and the current ratio $I_{Cl-}/I_{K+}$ are provided. Frames highlighted in orange (16 frames) exhibit a conductance less than 2.5 nS. These were used as starting point in MD simulations to better evaluate their conductance and stability.

| # | $\sigma$ | $I_{Cl-}/I_{K+}$ | # | $\sigma$ | $I_{Cl-}/I_{K+}$ | # | $\sigma$ | $I_{Cl-}/I_{K+}$ | # | $\sigma$ | $I_{Cl-}/I_{K+}$ |
|---|---|---|---|---|---|---|---|---|---|---|---|
| 1 | 1.20 | ¥ | 51 | 3.08 | 3.97 | 101 | 3.56 | 3.81 | 151 | 4.01 | 1.60 |
| 2 | 1.30 | 4.00 | 52 | 3.10 | 6.75 | 102 | 3.58 | 3.26 | 152 | 4.01 | 5.90 |
| 3 | 1.62 | 2.86 | 53 | 3.10 | 1.63 | 103 | 3.58 | 3.26 | 153 | 4.05 | 2.89 |
| 4 | 1.78 | 13.84 | 54 | 3.12 | 2.32 | 104 | 3.58 | 3.47 | 154 | 4.05 | 3.70 |
| 5 | 1.80 | 5.93 | 55 | 3.12 | 1.84 | 105 | 3.59 | 2.25 | 155 | 4.07 | 3.41 |
| 6 | 1.88 | 2.76 | 56 | 3.12 | 3.59 | 106 | 3.59 | 5.40 | 156 | 4.08 | 2.09 |
| 7 | 1.90 | 1.26 | 57 | 3.14 | 2.02 | 107 | 3.60 | 2.60 | 157 | 4.09 | 4.51 |
| 8 | 2.08 | 8.45 | 58 | 3.16 | 4.85 | 108 | 3.62 | 2.41 | 158 | 4.11 | 5.61 |
| 9 | 2.18 | 3.04 | 59 | 3.16 | 3.79 | 109 | 3.63 | 9.06 | 159 | 4.11 | 4.54 |
| 10 | 2.26 | 4.38 | 60 | 3.16 | 3.94 | 110 | 3.66 | 2.66 | 160 | 4.13 | 5.65 |
| 11 | 2.32 | 10.60 | 61 | 3.16 | 1.98 | 111 | 3.67 | 1.77 | 161 | 4.13 | 3.91 |
| 12 | 2.32 | 2.22 | 62 | 3.18 | 4.13 | 112 | 3.67 | 4.23 | 162 | 4.14 | 1.87 |
| 13 | 2.34 | 8.75 | 63 | 3.20 | 3.32 | 113 | 3.68 | 2.91 | 163 | 4.17 | 3.84 |

| | | | | | | | | | | | |
|---|---|---|---|---|---|---|---|---|---|---|---|
| 14 | 2.34 | 3.68 | 64 | 3.22 | 7.05 | 114 | 3.68 | 2.29 | 164 | 4.18 | 2.12 |
| 15 | 2.42 | 1.75 | 65 | 3.22 | 3.13 | 115 | 3.71 | 6.12 | 165 | 4.18 | 1.71 |
| 16 | 2.46 | 0.76 | 66 | 3.25 | 10.58 | 116 | 3.71 | 3.75 | 166 | 4.20 | 1.96 |
| 17 | 2.50 | 9.42 | 67 | 3.26 | 5.27 | 117 | 3.73 | 1.86 | 167 | 4.23 | 8.18 |
| 18 | 2.52 | 5.63 | 68 | 3.26 | 5.04 | 118 | 3.73 | 5.00 | 168 | 4.23 | 4.03 |
| 19 | 2.52 | 3.07 | 69 | 3.26 | 3.66 | 119 | 3.74 | 2.40 | 169 | 4.23 | 3.49 |
| 20 | 2.56 | 8.85 | 70 | 3.26 | 3.40 | 120 | 3.75 | 5.93 | 170 | 4.25 | 3.42 |
| 21 | 2.60 | 3.48 | 71 | 3.26 | 2.26 | 121 | 3.75 | 3.56 | 171 | 4.27 | 6.89 |
| 22 | 2.64 | 1.93 | 72 | 3.26 | 2.54 | 122 | 3.75 | 3.80 | 172 | 4.33 | 3.00 |
| 23 | 2.68 | 12.40 | 73 | 3.28 | 2.09 | 123 | 3.75 | 4.06 | 173 | 4.39 | 1.19 |
| 24 | 2.70 | 5.75 | 74 | 3.29 | 7.64 | 124 | 3.77 | 4.37 | 174 | 4.39 | 5.26 |
| 25 | 2.70 | 10.25 | 75 | 3.30 | 4.89 | 125 | 3.77 | 3.27 | 175 | 4.39 | 2.18 |
| 26 | 2.70 | 8.00 | 76 | 3.35 | 2.55 | 126 | 3.78 | 2.50 | 176 | 4.41 | 5.47 |
| 27 | 2.70 | 4.62 | 77 | 3.39 | 2.19 | 127 | 3.79 | 6.27 | 177 | 4.43 | 5.50 |
| 28 | 2.72 | 66.98 | 78 | 3.39 | 7.90 | 128 | 3.79 | 4.11 | 178 | 4.43 | 2.88 |
| 29 | 2.72 | 4.91 | 79 | 3.39 | 6.04 | 129 | 3.81 | 1.68 | 179 | 4.45 | 1.71 |
| 30 | 2.72 | 4.04 | 80 | 3.39 | 5.76 | 130 | 3.81 | 1.57 | 180 | 4.45 | 1.64 |
| 31 | 2.72 | 4.23 | 81 | 3.41 | 2.15 | 131 | 3.81 | 6.60 | 181 | 4.45 | 7.22 |
| 32 | 2.74 | 1.21 | 82 | 3.41 | 7.95 | 132 | 3.83 | 8.55 | 182 | 4.49 | 5.22 |
| 33 | 2.76 | 2.63 | 83 | 3.43 | 2.11 | 133 | 3.83 | 3.90 | 183 | 4.49 | 3.00 |
| 34 | 2.76 | 1.94 | 84 | 3.43 | 9.69 | 134 | 3.85 | 1.67 | 184 | 4.49 | 2.80 |
| 35 | 2.82 | 5.13 | 85 | 3.44 | 3.41 | 135 | 3.85 | 4.82 | 185 | 4.51 | 3.69 |
| 36 | 2.82 | 3.86 | 86 | 3.45 | 2.07 | 136 | 3.85 | 4.19 | 186 | 4.55 | 3.94 |
| 37 | 2.84 | 4.92 | 87 | 3.45 | 2.19 | 137 | 3.85 | 3.57 | 187 | 4.55 | 3.73 |
| 38 | 2.86 | 3.93 | 88 | 3.45 | 1.26 | 138 | 3.87 | 3.11 | 188 | 4.59 | 9.41 |
| 39 | 2.92 | 6.30 | 89 | 3.46 | 2.53 | 139 | 3.87 | 2.94 | 189 | 4.59 | 3.41 |
| 40 | 2.94 | 3.32 | 90 | 3.47 | 2.39 | 140 | 3.91 | 5.97 | 190 | 4.65 | 14.46 |
| 41 | 2.96 | 3.35 | 91 | 3.47 | 18.23 | 141 | 3.91 | 4.74 | 191 | 4.69 | 3.88 |
| 42 | 2.96 | 4.29 | 92 | 3.47 | 9.18 | 142 | 3.91 | 3.06 | 192 | 4.75 | 3.16 |
| 43 | 3.00 | 1.68 | 93 | 3.48 | 2.62 | 143 | 3.92 | 1.84 | 193 | 4.77 | 1.18 |
| 44 | 3.02 | 5.04 | 94 | 3.49 | 12.39 | 144 | 3.93 | 1.72 | 194 | 4.79 | 3.98 |
| 45 | 3.02 | 3.72 | 95 | 3.50 | 4.15 | 145 | 3.93 | 4.30 | 195 | 4.83 | 1.80 |
| 46 | 3.02 | 3.19 | 96 | 3.51 | 2.43 | 146 | 3.94 | 2.40 | 196 | 4.95 | 3.05 |
| 47 | 3.04 | 3.90 | 97 | 3.51 | 6.61 | 147 | 3.95 | 5.57 | 197 | 4.99 | 1.97 |
| 48 | 3.04 | 2.71 | 98 | 3.53 | 7.00 | 148 | 3.97 | 5.00 | 198 | 5.05 | 1.80 |
| 49 | 3.06 | 1.64 | 99 | 3.54 | 3.32 | 149 | 3.98 | 1.80 | 199 | 5.21 | 2.47 |
| 50 | 3.08 | 3.16 | 100 | 3.55 | 6.70 | 150 | 3.99 | 3.52 | 200 | 5.27 | 2.93 |
